# Dietary Analysis of *Vespula shidai* in Reared and Wild Nests

**DOI:** 10.1101/2024.08.18.608443

**Authors:** Tatsuya Saga

**Affiliations:** Graduate School of Human Development and Environment, Kobe University, 3-11 Tsurukabuto, Nada, Kobe, Hyogo, Japan 657-8501

## Abstract

This study examines the dietary behaviors of both reared and wild nests of *Vespula shidai* in Central Japan, a region known for its rich biodiversity and unique entomophagy culture, particularly the consumption of wasp larvae and pupae, known locally as “hachinoko.” This study reveal that these wasps exhibit remarkable dietary versatility, consuming a wide range of vertebrates—more varied than their counterparts in Hawaii and New Zealand. The study notes that wild nests, unlike their reared nests, consume a significantly greater variety of vertebrate species, even without supplementation. This suggests that vertebrate carcasses provide crucial nutrients that contribute substantially to wasp development. Additionally, this research highlights individual foraging preferences among nests, which are influenced by both the availability and nutritional value of prey, as well as human feeding practices. This variation in prey composition, particularly evident between reared and wild nests, underscores the complex interaction between wasps’ natural dietary habits and human intervention.

## Introduction

Japan’s rural woodlands and farmland, known as Satoyama, are high biodiversity areas and have sustained a unique food culture that includes entomophagy. In the mountainous regions of Central Japan, the larvae and pupae of social wasps such as *Vespula* and *Vespa* are consumed as “*hachinoko*” a term in Japanese for a delicacy of autumn (Nonaka 2009; Payne, 2015). The practices of collecting, rearing, and cooking these insects have continued to develop (Nonaka 2009; Payne and Evans 2017; Van Itterbeeck et al., 2021). Wasp enthusiasts not only cultivate wasp nests for consumption but also allow the next generation of queens to mate and hibernate, and release them back into nature in the spring (Nonaka 2009). Hachinoko eating represents a cuisine deeply rooted in the lifestyle and natural environment of the rural areas (Payne and Evans 2017). Additionally, these subfamily Vespinae wasps are not only valuable food sources but are also believed to suppress the populations of agricultural and sanitary pests (Ono 1997; Brock et al. 2021).

In Central Japan, when collecting nests of *Vespula shidai* for consumption, the wasps are given meat from chickens or squid formed into small balls with flags marked for tracking, which the workers carry back to their nests, allowing collectors to follow the flag to discover the nests (Saga 2019). Local enthusiasts reare these wasps using a variety of foods, including chicken, frog, deer meat, and fish. They select the feed based on their experience and the amount consumed by the reared workers, yet they have limited knowledge of other species the workers might consume beyond the provided feed. Although 54 prey species for *Vespula* wasps in Japan have been documented by Shida (1963) and Iwata (1971), there has been little additional knowledge since then.

Recently, the use of genetic methods to identify the prey species of social hunting wasps has been proposed (Takahashi et al., 2016). Furthermore, the DNA metabarcoding technique, which involves decoding the nucleotide sequences from fragments of prey organisms found in larval digestive tracts or left in feces in the nests using high-throughput sequencing, has made it possible to identify a large number of prey species at once (Lefort et al., 2020; Howse et al., 2022).

Wasps in the subfamily Vespinae, including *Vespula* and *Vespa*, retain feces in their digestive tracts during the larval stage and excrete them all at once when entering the prepupal stage. When becoming pre-pupae, feces are expelled at the bottom of the brood cell (Spradbery 1973). This study aims to use this method to identify the prey species from the contents of *V. shidai* larvae’s digestive tracts.

## Materials and Methods

### 1. Life History and Characteristics of *Vespula shidai*

The nesting behavior of *V. shidai* resembles that of other *Vespula* species (Spradbery 1973; Matsuura 1995). Each spring, a single queen emerges from hibernation and starts a new colony. After hibernation, the queen establishes the colony and starts producing workers in April or May. Early in the season, *V. shidai* queens often take over the nests of other queens from their own species, as well as those of their sister species, *Vespula flaviceps* (Saga et al. 2017). The colony grows throughout the summer and engages in reproductive activities from autumn to winter. In the fall, the queen mates with several males. By the beginning of winter (around November in central Japan), only the new queens go into hibernation while the workers and males die (Matsuura and Yamane 1990; Saga et al. 2020).

### 2. Sampling

In this study, we collected five wild nests in Tsukechi town and Takayama city, and assigned the care of seven reared nests to specific local enthusiasts in Tsukechi town, Ena city, Kushihara town in Gifu prefecture, and Anan town in Nagano Prefecture. I baited wild nests with Japanese dace and chicken meat. Each town or city had a designated enthusiast responsible for the nests. From each nest, we used 1 to 6 final-stage larvae for dietary analysis. In Tsukechi town, the appointed enthusiast fed captive nests with chicken, quail, deer, and wild boar meat, while in Ena City and Kushihara, only chicken was provided, and no feed was given in Anan Town. The reared nests in Tsukechi Town were randomly selected each year from the same site where 20 nests were maintained. Additionally, the wild nests in Tsukechi Town were collected from the same mountain slope within a 5 km radius of the rearing site. Table 1 summarizes the timing of nest collection (and sample collection for dietary analysis from captive nests), as well as the number of nests and individuals used. I collected undigested material from the intestines using the same method described by Takahashi et al. (2016). I collected approximately 2mm^3^ of gut content from the mouth-side of the intestine.

**Table 1.**
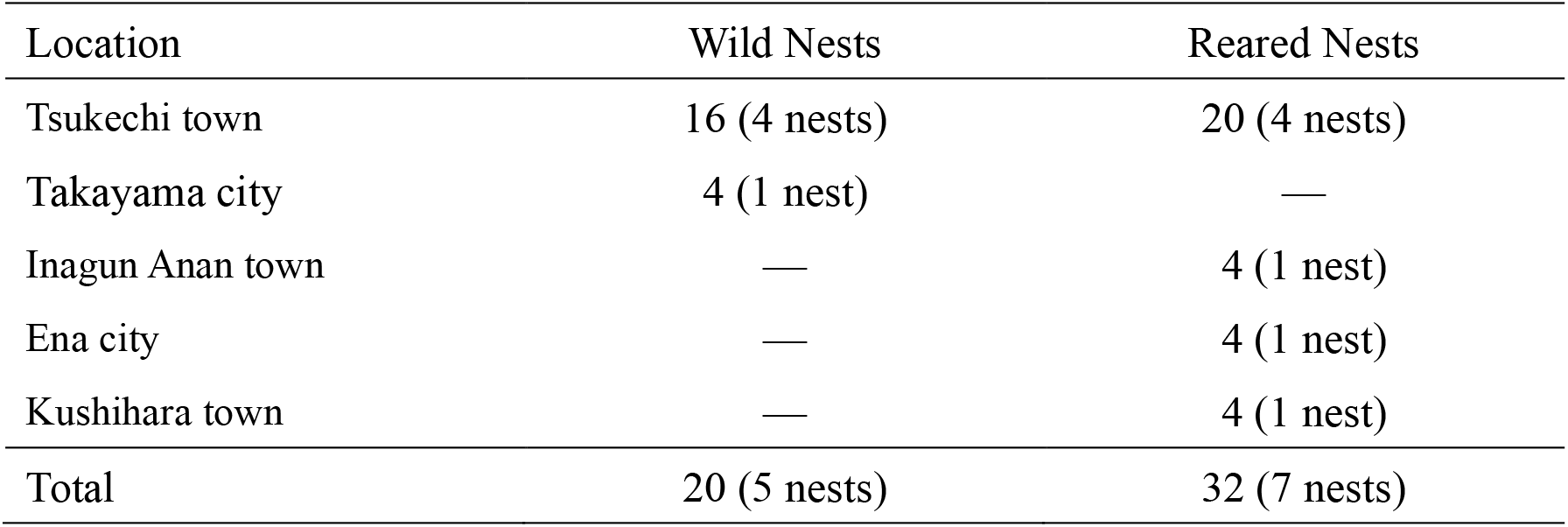
Number of worker larvae used for feed analysis.

### 3. DNA Experiments

#### (1) DNA Extraction and Amplification

I extracted genomic DNA using the NucleoSpin® Tissue kit (MACHEREY– NAGEL, Germany) following standard procedures.

#### (2) 1st PCR

I used 1st-IntF (Leray et. al 2013) and 1st-HCOmR (Folmer et. al 1994) primers to amplify the mitochondrial DNA COI region. I performed the PCR with TaKaRa Ex Taq Hot Start Version (Takara Bio Inc., Japan). To prevent amplification of the insect’s own DNA from intestinal prey, I used a blocking primer specifically for V. shidai. The PCR conditions were an initial 2-minute reaction at 94°C, followed by 35 cycles of 30 seconds at 94°C, 15 seconds at 67°C, 30 seconds at 52°C, 30 seconds at 72°C, and a final extension for 5 minutes at 72°C.

#### (3) 2nd PCR

I performed tailed PCR using 2nd primers that include index sequences and KOD FX Neo (TOYOBO) at 0.20uL (1.0 U/uL). The PCR conditions started with a 2-minute reaction at 94°C, followed by 12 cycles of 10 seconds at 98°C, 30 seconds at 60°C, 30 seconds at 68°C, and a final 2-minute extension at 68°C. After the PCR, we added VAHTS DNA Clean Beads (Vazyme) in an amount equal to the PCR volume and purified the PCR products.

#### (4) Sequencing

I sequenced using the MiSeq system and MiSeq Reagent Kit v3 (Illumina) under 2×300 bp conditions. We used Qiime2 (ver. 2020.8) with the dada2 plugin to remove chimera and noise sequences, then created representative sequences and Operational Taxonomic Units (OTUs). I conducted BLASTN (ver.2.13.0) for phylogenetic estimation and compared sequences with those in Gen Bank (National Center for Biotechnology Information, USA) to identify species.

#### (5) Dilution

I removed low-frequency OTUs with fewer than 5 reads. I then used the rrarefy function in R to resample from each sample according to the obtained read counts, normalizing the variation in total read counts of OTUs obtained from each wasp larva.

#### (6) Species identification

I identified sequences as belonging to the same species as the top-ranked reference in the GenBank database if the sequence similarity was 97% or higher, based on bit scores. If different species showed the same sequence similarity but different bit scores, I did not specify the species and instead considered it as belonging to the same genus, family, or order. I considered sequences with less than 97% similarity as unidentified. I also removed sequences of non-target species, the wasp themselves, mites, and humans due to the inability to distinguish them from contaminants.

### 4. Statistical Analysis

I conducted a t-test to see if there were differences in the number of prey species between reared and wild nests of *V. shidai*. Additionally, to compare the prey species composition of each larva, I calculated the dissimilarity using the Bray-Curtis index and applied non-metric multidimensional scaling (NMDS). I then conducted PERMANOVA (permutational analysis of variance), using nest type (reared or wild) and regional differences as factors, to test for differences in prey species. After identifying regional differences in prey composition, I specifically analyzed the dissimilarity of larval diets in nests from Tsukechi Town, where comprehensive data from both reared and wild nests were available. Then I conducted PERMANOVA to see if there were differences in prey composition between reared and wild nests. All statistical analyses were conducted using R (R Core Team 2023).

## Results

### Diet of reared and wild nests of *Vespula shidai*

We analyzed the gut contents of *Vespula shidai* larvae from 12 nests, identifying 324 prey species across 119 families and 29 orders. The orders with significant diversity included Lepidoptera (163 species), Diptera (39 species), Hemiptera (27 species), Araneae (22 species), Hymenoptera (13 species), Orthoptera (9 species), Coleoptera (8 species), Psocodea (6 species), Odonata (4 species), and other insects/arachnids (20 species). Additionally, I identified 10 bird species and 13 other vertebrate species (including fish, reptiles, amphibians, and mammals).

In the study, several species of vertebrates were detected across different categories: Birds: The species detected included the Japanese quail (*Coturnix japonica*), chicken (*Gallus gallus domesticus*), Japanese bush warbler (*Horornis diphone*), Eurasian jay (*Garrulus glandarius*), Large-billed crow (*Corvus macrorhynchos*), Carrion crow (*Corvus corone*), warbling white-eye (*Zosterops japonicu*s), Kamchatka leaf warbler (*Phylloscopus examinandus*), brown-eared bulbul (*Hypsipetes amaurotis*), and the oriental Turtle Dove (*Streptopelia orientalis*). Mammals: Among mammals, the wild boar (Sus scrofa), Sika deer (*Cervus nippon*), masked palm civet (*Paguma larvata*), Japanese weasel (*Mustela itatsi*), Japanese serow (*Capricornis crispus*), and Japanese grass vole (*Microtus montebelli*) were detected. Amphibians: Detected amphibians included Schlegel’s green tree frog (*Rhacophorus schlegelii*), forest green tree frog (*Rhacophorus arboreus*), and Japanese tree frog (*Hyla japonica*). Reptiles: The Japanese grass lizard (*Takydromus tachydromoides*), Eastern Japanese common skink (*Plestiodon finitimus*), and Japanese forest gecko (*Gekko japonicus*) were found. Fish: The study also noted the presence of the Japanese dace (*Tribolodon hakonensis*).

The average number of prey species per nest was 48.4 ± 34.89 (SD, n = 5) in wild nests and 45.6 ± 19.88 (n = 7) in reared nests, with no significant difference between them (t-test, p = 0.874). Vertebrates specifically showed a differential pattern, with wild nests containing an average of 5.0 ± 1.10 species and reared nests 3.71 ± 1.39. Notably, wild nests had significantly more wild vertebrate species (4.0 ± 1.55) compared to reared nests (1.71 ± 1.16) (t-test, p = 0.023).

Bird DNA was detected in all examined nests. Mammalian DNA was found in all but two nests—one wild nest in Tsukechi and one reared nest in Anan Town—indicating broad consumption patterns. The other ten nests showed evidence of mammalian predation. In Tsukechi Town, one reared nest, intentionally fed with deer, tested positive for deer DNA. Deer DNA was also found in a wild nest in Tsukechi Town and another reared nest in Kushihara Town, neither of which were fed deer.

I illustrated the composition of prey species for each collection region of *V. shidai* (stress value = 0.251, n = 44, Figure 1). Significant differences were observed in prey species similarity between reared and wild nests, and across collection/breeding regions (PERMANOVA; n = 44; reared or wild: R2 = 0.08, p < 0.01; collection region: R2 = 0.162, p < 0.01; Nest Differences: R2 = 0.05, P < 0.01). Further analysis of nests from Tsukechi Town, where both reared and wild data were available, revealed distinct differences in prey composition based on nest type (PERMANOVA; n = 36; reared or wild: R2 = 0.13, P < 0.01; Nest Differences: R2 = 0.340 P < 0.01; Year: R2 = 0.08, P < 0.01, Figure 2).

**Figure 1.**
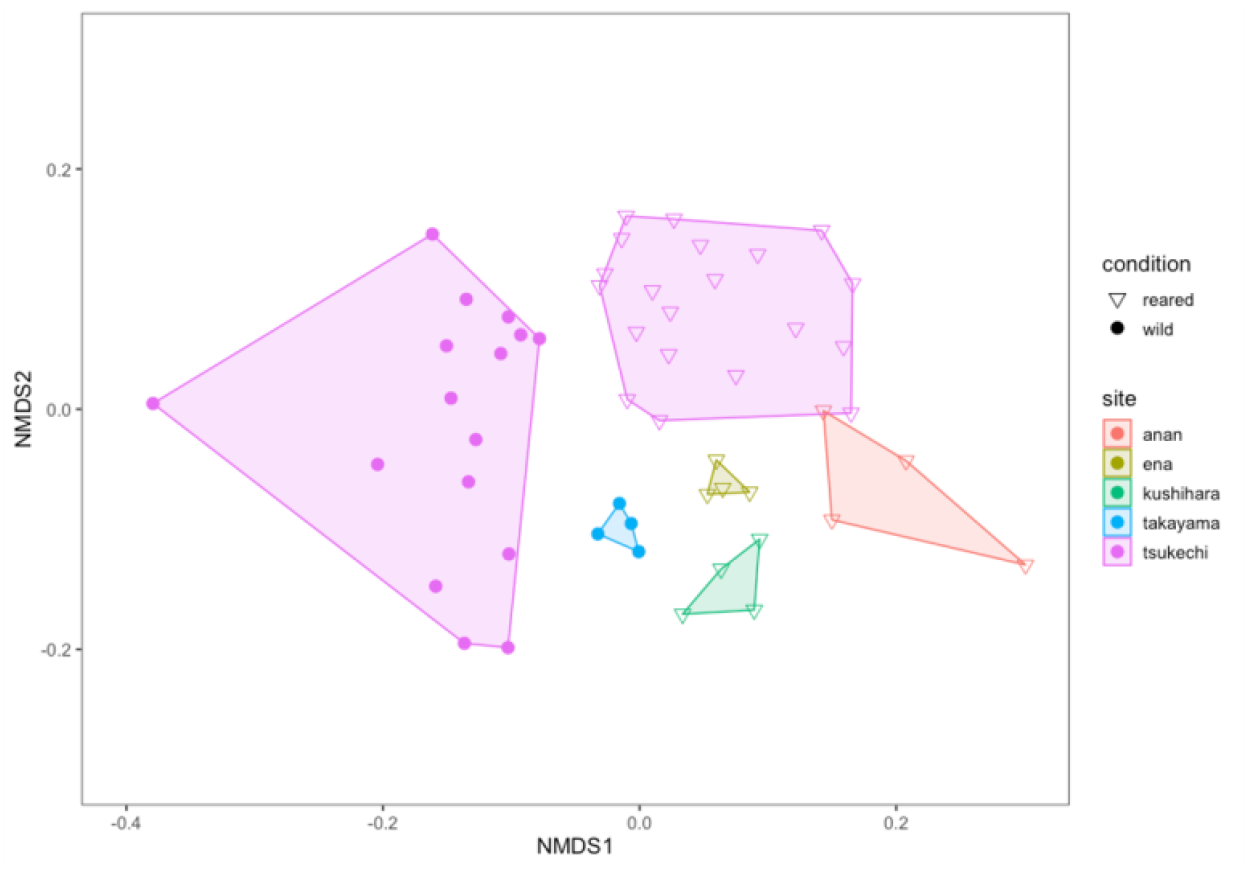
Similarity in diet composition between reared and wild nests of *Vespu/a shidaiin* five municipalities (created using the Bray-Curtis index, stress value= 0.251, n = 44). Significant differences in the similarity of prey species were observed depending on whether the nests were reared or wild, and by the region where the wasps were collected and reared (PERMANOVA; n = 44; reared or wild: R^2^ = 0.08, p < 0.01; collection region: R^2^ = 0.162, p < 0.01; nest differences: R^2^ = 0.05, p < 0.01).

**Figure 2.**
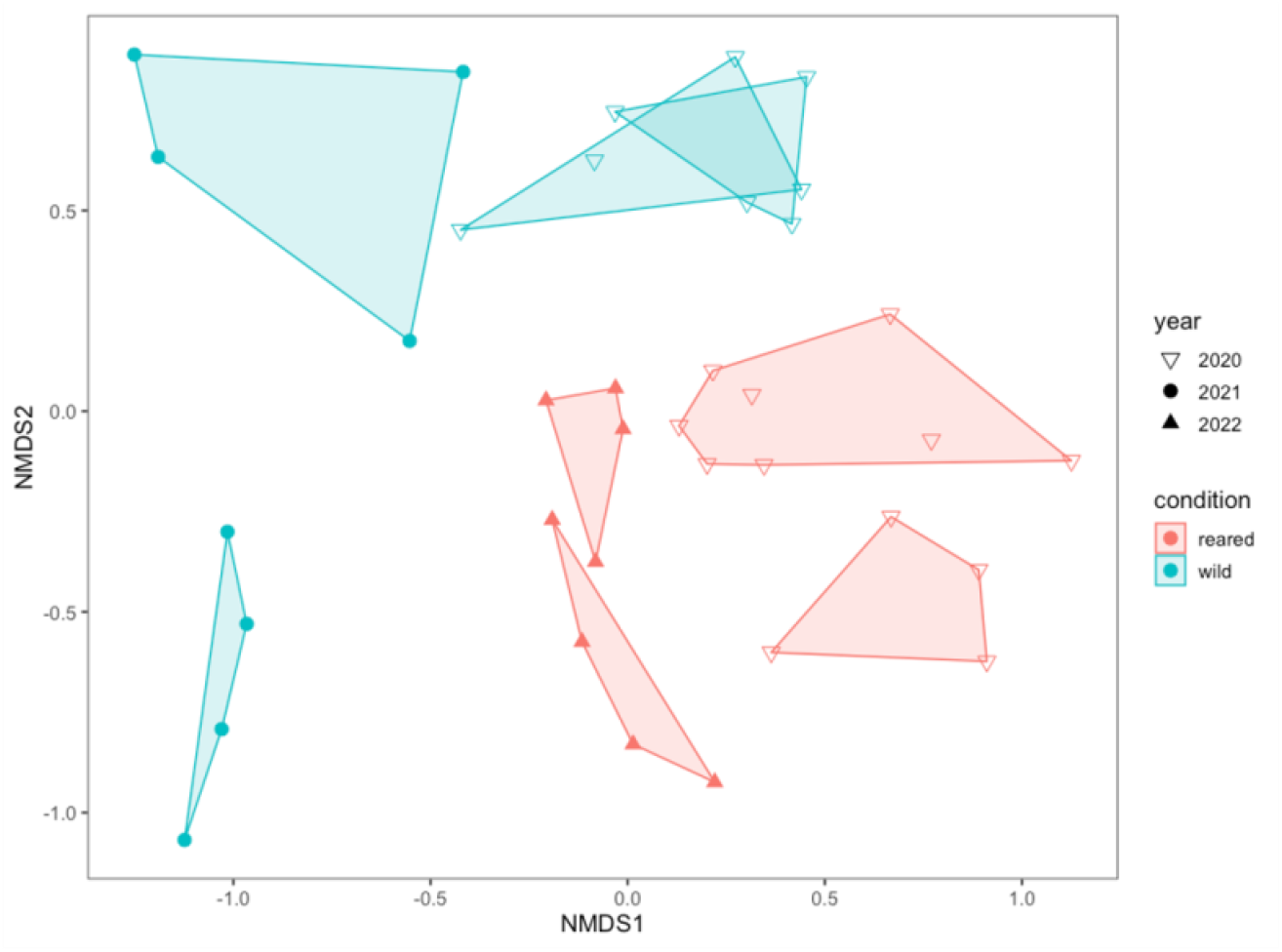
Similarity in diet composition between captive and wild nests of *Vespula shidaiin* Tsukechi town. Differences in prey composition were observed depending on whether the nests were reared or wild, and by the specific nest (PERMANOVA; n = 36; reared or wild: R^2^ = 0.13, p < 0.01; nest differences: R^2^ = 0.340, p < 0.01; Year: R^2^ = 0.08, p < 0.01).

## Discussion

This study reveals that *Vespula shidai* exhibits a diverse dietary range, consuming the most extensive variety of prey species among vertebrates, observed in both reared and wild nests. This finding indicates a higher level of dietary versatility compared to *Vespula* species studied in Hawaii and New Zealand. The detection of a wide array of prey—from birds to mammals, frogs and geckos—underscores the rich biodiversity of Central Japan’s regions and the adaptive dietary strategies of *V. shidai*.

### Comparative analysis with global counterparts

In Hawaii, workers from ten nests of the invasive *Vespula pensylvanica* were found to bring back pieces of meat that, when analyzed using DNA barcoding, revealed a prey spectrum comprising 47 families across 14 orders, including vertebrates such as pheasants (Phasianidae), perching birds (Passeriformes), rodents (Muridae), and geckos (Gekkonidae) (Wilson et al. 2009). In contrast, on Ahuahu Island in New Zealand, the application of DNA metabarcoding to nest feces identified 33 prey species including rodents from three nests (9 individuals) of *Vespula germanica*, and 68 prey species including shorebirds from 16 nests (48 individuals) of *Vespula vulgaris* (Schmack et al. 2021). *V. shidai* in my study displayed the broadest prey spectrum, evidenced by the detection of 324 prey species across 119 families and 29 orders from just twelve nests, indicative of the rich prey availability and possibly the intricate feeding behavior fostered by the diverse ecosystems of Central Japan’s rural woodlands.

### Local rearing practices and their ecological implications

This research has also confirmed that reared nests, where wasps are fed human-provided meats such as chicken and deer, indeed consume these foods. However, it was notable that wild nests also preyed on bird and mammal meats, which had not been previously well-documented. This observation not only confirms the sophistication of local rearing practices but also suggests that information exchange among breeders about suitable meats has likely influenced feeding practices. The results emphasize that *V. shidai* is capable of utilizing a variety of vertebrates as part of their natural diet, which aligns with the observed preferences in different nests.

### Dietary diversity and foraging strategies

This study highlights significant dietary behaviors of both reared and wild *V. shidai* nests, noting that birds and mammals are common prey across almost all nests. Particularly, wild nests that were not supplemented with vertebrate prey captured a significantly greater variety of vertebrate species than reared nests, suggesting that vertebrate carcasses, generally larger than arthropods and potentially rich in essential nutrients, contribute substantially to wasp development. The findings challenge the notion that wasps merely gather available prey near their nests, as it was observed that nests reared at the same site displayed varied prey compositions. This suggests that individual nests may have distinct foraging preferences, influenced by the availability and nutritional value of prey. The study also notes that the type of feed provided by enthusiasts could alter natural foraging behaviors, indicating that human management practices have a tangible impact on wasp feeding strategies and, by extension, their ecological roles.

The disparities in prey composition between reared and wild nests of *V. shidai* reveal the complexity of their dietary habits, influenced by both local environmental conditions and human intervention. This variation is crucial for understanding their survival, reproductive success, and the broader ecological roles these wasps play. These findings emphasize the need for species-specific feeding strategies and underscore the importance of incorporating both natural and anthropogenic factors into conservation and management practices to ensure the sustainability of wasp populations and Hachinoko food culture.

## Conclusion

In summary, the refined breeding culture and the varied dietary intake of *V. shidai* highlight the adaptability and ecological impact of this species in Central Japan, offering insights into their potential role in local biodiversity management and conservation strategies.

## Acknowledgments

I am deeply grateful for the support I received during this research. Part of the experiments was conducted with the cooperation of Mr. Hiroyasu Kinoshita. Dr. Junichi Takahashi provided valuable advice on experimental methods. I would like to express my deepest gratitude here. The wasp nests were provided by Mr. Katsuyuki Takahashi, Mr. Fumihiro Sato, and Mr. Osamu Fukatsu. I am thankful for their efforts in collecting these wasps. This research was partially supported by grants from the Asahi Group Foundation for Academic Promotion, Takara Harmonist Fund, Nissei Foundation Environmental Research Grant, Takeda Science Foundation Grant for the Promotion of Science Education in High Schools, Ito Youth Scholarship Association Regional Promotion Grant, JSPS KAKENHI Grant Number 21H04158, Shimonaka Memorial Foundation, and TAKEO Co., Ltd. I extend my gratitude to all for their support.

## References

Brock, R. E., Cini, A., & Sumner, S. (2021). Ecosystem services provided by aculeate wasps. Biological Reviews, 96(4), 1645–1675.

Folmer, O., Black, M., Hoeh, W., Lutz, R., & Vrijenhoek, R. (1994). DNA primers for amplification of mitochondrial cytochrome C oxidase subunit I from diverse metazoan invertebrates. Molecular Marine Biology and Biotechnology, 3, 294–299.

Howse, M. W. F., McGruddy, R. A., Felden, A., et al. (2022). The native and exotic prey community of two invasive paper wasps (Hymenoptera: Vespidae) in New Zealand as determined by DNA barcoding. Biol Invasions, 24, 1797–1808. 10.1007/s10530-022-02739-0

Iwata, K. (1971). Honnou no Shinka: Hachi no Hikaku Shuseigakuteki Kenkyu. Mano Shoten.

Leray, M., Yang, J. Y., Meyer, C. P., Mills, S. C., Agudelo, N., Ranwez, V., Boehm J. T., & Machida, R. J. (2013). A new versatile primer set targeting a short fragment of the mitochondrial COI region for metabarcoding metazoan diversity: application for characterizing coral reef fish gut contents. Frontiers in Zoology, 10(1), 1–14.

Nonaka, K. (2009). Feasting on insects. Entomological Research, 39(5), 304–312.

Ono, M. (1997). Suzumebachi no Kagaku. Kaiyusha.

Payne, C. L. R. (2015). Wild harvesting declines as pesticides and imports rise: the collection and consumption of insects in contemporary rural Japan. Journal of Insects as Food and Feed, 1(1), 57–65.

Payne, C. L. R., & Evans, J. D. (2017). Nested Houses: Domestication dynamics of human–wasp relations in contemporary rural Japan. Journal of Ethnobiology and Ethnomedicine, 13, 13. 10.1186/s13002-017-0138-y

Schmack, J. M., Lear, G., Astudillo-Garcia, C., Boyer, S., Ward, D. F., & Beggs, J. R. (2021). DNA metabarcoding of prey reveals spatial, temporal and diet partitioning of an island ecosystem by four invasive wasps. Journal of Applied Ecology, 58(6), 1199–1211.

Shida, T. (1963). Tokyo Kinko ni Okeru Kurosuzumebachi no Emono. Konchu, 31, 198–199.

Spradbery, J. P. (1973). Wasps: An Account of the Biology and Natural History of Solitary and Social Wasps. Sidgwick & Jackson, London.

Takahashi, R, Kiyoshi T, & Takahashi J. (2016). An attempt to identify the diets of Vespa velutina using the DNA barcoding method. Nagasaki-ken Seibutsugaku Kai Shi, (78), 43–48.

Van Itterbeeck, J., Feng, Y., Zhao, M., Wang, C., Tan, K., Saga, T., … & Jung, C. (2021). Rearing techniques for hornets with emphasis on Vespa velutina (Hymenoptera: Vespidae): A review. Journal of Asia-Pacific Entomology, 24(2), 103–117.

Wilson, E. E., Mullen, L. M., & Holway, D. A. (2009). Life history plasticity magnifies the ecological effects of a social wasp invasion. Proceedings of the National Academy of Sciences, 106(31), 12809–12813.

